# Domain prediction with probabilistic directional context

**DOI:** 10.1101/094284

**Authors:** Alejandro Ochoa, Mona Singh

## Abstract

**Motivation:** Protein domain prediction is one of the most powerful approaches for sequence-based function prediction. While domain instances are typically predicted independently of each other, newer approaches have demonstrated improved performance by rewarding domain pairs that frequently co-occur within sequences. However, most of these approaches have ignored the order in which domains preferentially co-occur and have also not modeled domain co-occurrence probabilistically.

**Results:** We introduce a probabilistic approach for domain prediction that models “directional” domain context. Our method is the first to score all domain pairs within a sequence while taking their order into account, even for non-sequential domains. We show that our approach extends a previous Markov model-based approach to additionally score all pairwise terms, and that it can be interpreted within the context of Markov random fields. We formulate our underlying combinatorial optimization problem as an integer linear program, and demonstrate that it can be solved quickly in practice. Finally, we perform extensive evaluation of domain context methods and demonstrate that incorporating context increases the number of domain predictions by ∼15%, with our approach dPUC2 (Domain Prediction Using Context) outperforming all competing approaches.

**Availability:** dPUC2 is available at http://github.com/alexviiia/dpuc2.

## 1 Introduction

Protein domains are the functional and evolutionary units of proteins; they organize protein space and their identification helps annotate new protein sequences. Domains are often predicted in protein sequences using profile Hidden Markov Models (HMMs) [1], each of which models one domain family and is constructed from sequence alignments of known instances. In the standard approach, statistical significance is evaluated for every domain prediction independently; this is how domains are identified in Pfam, the largest HMM domain database [2], by the state-of-the-art HMMER3 program [1]. Several recent methods have proposed to evaluate domains in the context of other domain predictions, using higher-order scoring systems and additional information; these include a Markov model of sequential domain co-occurrence [3], a domain prediction filter based on pairwise co-occurrence (CODD) [4], and a multi-objective optimization that uses known domain architectures (DAMA) [5]. Our previous approach, dPUC (Domain Prediction Using Context), maximizes the log-odds score of domain predictions and incorporates pairwise co-occurrence probabilities [6]. These earlier approaches have demonstrated that context yields improvement in domain identification, but have only been evaluated in small tests, with a focus on the compositionally-biased *Plasmodium falciparum* proteome [4–7].

It is well-known that domains tend to have order preferences within protein sequences [8]; in Pfam 25 [2], we observe that 88.59% of domain family pairs occur in only one orientation (i.e., where a domain instance of one family is always found earlier in a protein sequence than a domain instance of the other family). However, previous context-based approaches either do not consider this directionality [4–6] or only consider it for adjacent domain pairs [3]. In this work, we extend our earlier log-odds model for context-based domain prediction [6] to use directional context scores. We also newly show that our approach can be equivalently viewed as a Markov random field, and explain its connection to the earlier Markov model of [3]. We introduce a new integer linear programming (ILP) formulation to solve the underlying optimization problem, and demonstrate that in practice solutions can be obtained fast enough for routine application. Lastly, we demonstrate superior performance of our approach, dPUC2, over all of its publicly available competitors in testing based on the UniProt database [9]; this is the largest-scale testing to date of context-based domain prediction. Overall, we find that the benefits of using domain context is substantial, with ∼15% more domains predicted by dPUC2 than the Standard Pfam at the same level of estimated accuracy.

## 2 Models

### 2.1 The basic probabilistic model

Given a set of candidate domains, we develop an approach that identifies the most likely subset of domains under a probabilistic domain co-occurrence model. Let *𝓓* = {*D*_1_,*D*_2_, …, *D_k_*} be *k* candidate domain instances ordered by start coordinates, which are predicted on a protein sequence *X* with a permissive p-value threshold (e.g., using HMMER3). Let *F_i_* be the domain family of the domain instance *D_i_*; note that domain instances are individual predictions, while domain families correspond to Pfam HMMs. Our goal is to identify *𝓓′ ⊆𝓓* that maximize a score consisting of two log-odds components: an individual domain score and a pairwise domain context score. We additionally require that all domain instances *𝓓′* are compatible with each other. In particular, we do not allow predicted domain instances to overlap by > 40 amino acids or > 50% of the smaller of the two ranges; this is the “permissive overlap” definition used earlier [10, 11].

The individual domain scores are derived from HMMER, which scores domain instance *D_i_* as

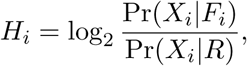

where *X_i_* is the subsequence of the predicted domain, Pr(*X_i_*|*F_i_*) is the probability of *X_i_* given the HMM of *F_i_* (the alternative model), and Pr(*X_i_*|*R*) is the probability of *X_i_* under the null model *R* of independent and identically distributed amino acids [1].

Pfam provides expert-curated “domain” thresholds 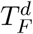 for each family *F*, where in order to predict a domain instance *D_i_* of family *F_i_*, it must have a score 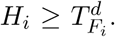 These thresholds have been previously interpreted as incorporating a prior probability [3]; in order to better fit a log-odds framework, our interpretation differs slightly by including Pr(*R*). Here, if Pr(*F_i_*) and Pr(*R*) are prior probabilities for domain family *F_i_* and the background model*R*, respectively, then the domain threshold is viewed as

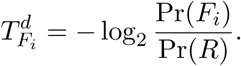

In this case,

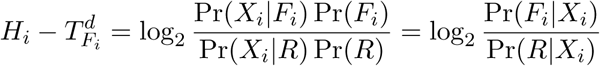

corresponds to a posterior log-odds score.

In addition to individual domains, we also incorporate “context” scores between domain families
as

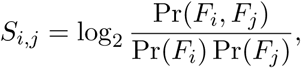

where Pr(*F_i_*, *F_j_*) is the probability that a domain of family *F_i_* appears earlier than one of family *F_j_* in the protein sequence, and Pr(F*_i_*) is the prior probability of observing domains of *F_i_*.

Our goal is to find *𝓓′* to maximize

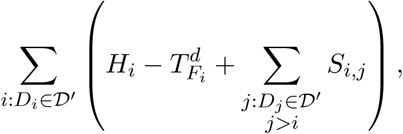

which is equivalent to

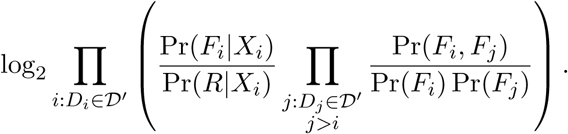

Hence, our objective is a log-odds score with two parts: the fit of each family to the sequence, and of family pairs to the co-occurrence model. Each part is a ratio of corresponding probabilities from null and alternative models. Note that no domains are predicted if the maximum value of the objective function is less than 0, since in this case the null model has the highest probability.

### 2.2 Formulation as a Markov random field

We note that we can also frame our model as a Markov Random Field (MRF). A configuration x = (*x*_1_,…, *x_k_*) in the system consists of indicator variables *x_i_* ∊ {0,1} for whether domain instance *i* is predicted or not. Let *𝓧* be the set of configurations without conflicting domains. The probability of x ∊ *𝓧* is

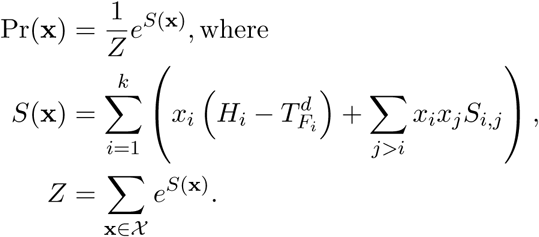

The solution x ∊ *𝓧* that maximizes Pr(x) is equivalent to the one that maximizes the log-odds score outlined above as *Z* is a normalization constant shared by all x.

### 2.3 Comparison to the Markov model of Coin *et al*.

Previous work [3] derived a Markov approximation for the joint probability of domain predictions given the sequence and the model. Using our notation and ignoring transitions from the “begin” and to the “end” states, for a given set of domains, *D*_1_… *D_k_*, the first-order Markov score is

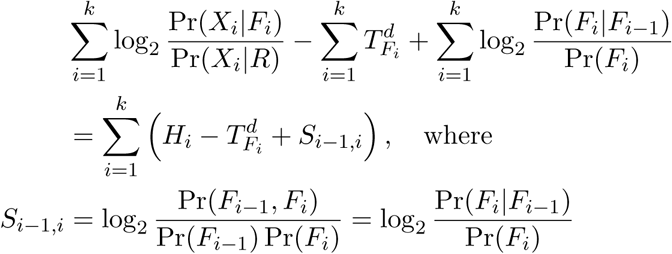

is our context score for consecutive domains (*S*_0,1_ =0 is set for convenience). Hence, our objective differs from the Markov model of [3] by

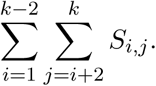

Notably, both scores agree for *k* ≤ 2 domain predictions, and for *k* = 3 the difference is only *S*_1,3_. Our approach has the advantage over the Markov model of enforcing consistency between all domain pairs.

### 2.4 Modifications due to Pfam thresholds

In addition to domain thresholds 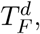 the Standard Pfam approach also uses expert-curated “sequence” thresholds 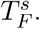 Let *𝓓*′*_F_* denote the subset of predictions within a protein that are domain instances of family *F*. These predicted domain instances must satisfy

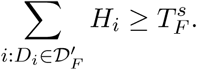

For 97.7% of Pfam families [12],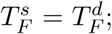 in other words, the sequence thresholds can be ignored since by satisfying the individual domain thresholds, the sequence thresholds are also satisfied. However, in order to make our model consistent with Pfam for the remaining 2.3% of Pfam families, we modify our model to include the term

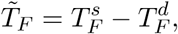

despite the fact that these sequence thresholds do not have a probabilistic interpretation. Our objective then becomes to find *𝓓*′ that maximize

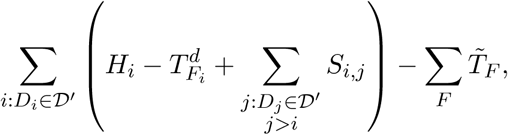

where every predicted family is included once in the sum over *F*.

Like previous models [3, 6], our context-based approach is designed to agree with the Standard Pfam in the absence of context (all *S_i_,_j_ =* 0), where the objective becomes

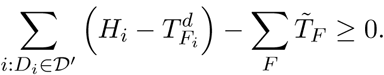

Each *D_i_* in the optimal solution satisfies 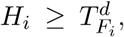 since otherwise including *D_i_* would reduce the objective. By the same reasoning, each family *F* in the optimal solution satisfies its sequence threshold, since

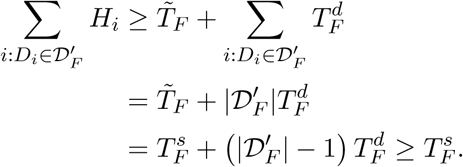

The sequence threshold is set exactly for 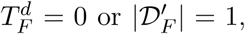 which are the most common cases. Otherwise 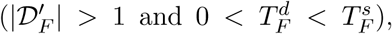 our approach’s effective sequence threshold is more stringent than the Standard Pfam’s, since the last inequality above becomes a strict inequality.

## 3 Methods

### 3.1 Context score estimation

We use a directed “context network” of family pair counts *c_i_,_j_* (≠ *C_j_,_i_*) of domain instances of family *F_i_* that occurred before instances of family *F_j_* in a large database. Self pairs (*F_i_* = *F_j_*) model repeating domains. Pair probabilities, corresponding to the proportion of all domain pairs that are of a given family pair, are estimated using a symmetric Dirichlet prior with parameter *α*, yielding

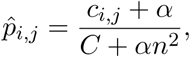

where *n* is the number of domain families (there are *n*^2^ directed family pairs, including self pairs) and 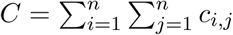 sums over family pairs. All families have equal prior probabilities 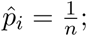 we note that the Bayesian interpretation of Pfam thresholds outlined earlier would set different values of Pr(*F_i_*), but these equal priors are simple and work well in practice. These estimates satisfy 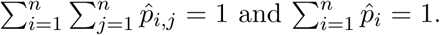 The context score for a domain instance of family *F_i_* preceding an instance of family *F_j_* is estimated as

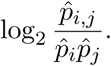

Note that unobserved family pairs (c_*i,j*_ = 0) receive a fixed penalty of

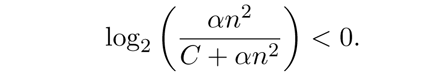

Counts *c_i_,_j_*are obtained from domains predicted by Pfam on UniProt, as provided in the file Pfam-A.full. We remove *c_i_,_j_* = 1 cases at load time, which are assumed to be false positives. We set *α =* 10^−3^, which we find to perform better than the uniform Dirichlet prior where *α* = 1.

### 3.2 Algorithms

We find optimal domain architectures with two strategies: “positive elimination” (PE), followed by Integer Linear Programming (ILP). PE discards domains with poor HMMER3 and context scores, returning trivial solutions (see below) or passing a smaller set to the ILP solver. The combinatorial ILP problem is solved using the lp_solve C library [13]; though ILP is NP-complete, in practice our optimization problem is solved quickly compared to the time taken for predicting the initial candidate domains using a permissive threshold via HMMER3 (see Section 4.2 below).

### 3.2.1 Positive elimination (PE)

PE consists of the following iterative procedure [6]. At a given iteration, remaining domain instances *D_i_* are evaluated using

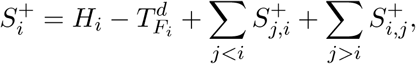

where only positive context scores 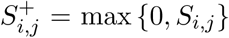 are included between *D_i_*and all remaining *D_j_*, and additionally 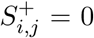for conflicting domains *D_i_*,*D_j_*. Domain *D_i_* is eliminated if 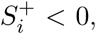 and all instances of family *F* are eliminated if 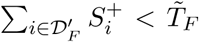. Note that eliminated *D_i_* would contribute negatively to our objective, even when every positive context score is included. Iterations end when there are no more eliminations.

After PE, the remaining domains are tested for conflicts or negative context: if both are absent, then these domains are the trivial solution and are returned directly, skipping the ILP. This follows since all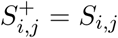 in this case, so 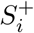 is the total score contributed by *D_i_*. Since every remaining *D_i_* satisfies 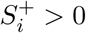 and 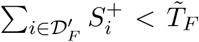, then all these *D_i_* improve the score, so they are all in the optimal solution.

For *k* domains, one PE iteration and the “trivial solution” test run in *O*(*k*^2^).

### 3.2.2 Integer Linear Program (ILP)

Let *x_i_*,*x_i_,_j_*, and *x_F_* be indicator variables for whether domain instances *D_i_*, pairs *D_i_* and *D_j_*, and families *F* are predicted, respectively. The objective function is

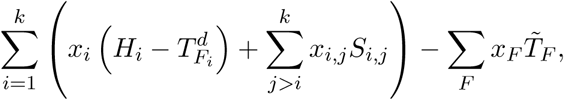

where *k* is the number of candidate domains and the sum over *F* covers all candidate families. We wish to find values for all *x_i_*, *x_i_,_j_*, and *x_F_* such that

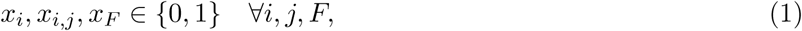

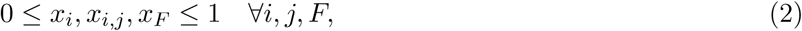

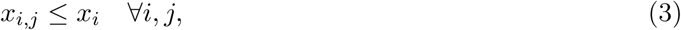

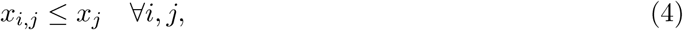

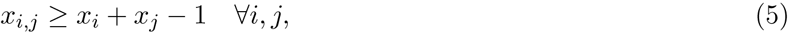

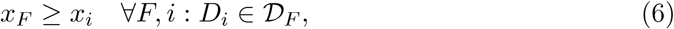

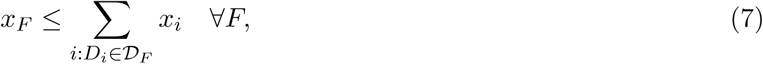

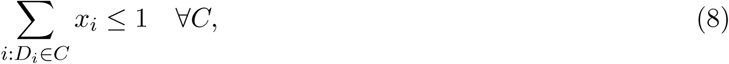

where *𝓓_F_* is the set of candidate domains of family *F*, and each *C* is a set of domains where all pairs conflict, and such that all conflicts among all possible domain instances are covered by all *C*’s. Note that the LP relaxation of this ILP formulation corresponds to dropping Eq.(1), and that constraint (2) is implied by constraint (1) in the ILP.

In the ILP, constraints (3)–(5) correspond to the “AND” operation of *x_i_,_j_* = *x_i_* Λ *x_j_*, where the pairwise domain variable *x_i_,_j_* is set to 1 if and only if both domains *D_i_* and *D_j_* are chosen. This is the tightest set of constraints implementing this requirement, as the four allowed settings of these variables are vertices on the resulting polytope (Fig. 1A). In contrast, Fig. 1B shows that an earlier formulation of the “AND” constraint [6] is not as effective. Constraints (6)–(7) in the ILP formulation correspond to the “OR” operation of 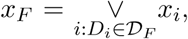 where the family variable *x_F_* is set to 1 only if at least one domain instance of that family is chosen. Lastly, constraint (8) in the ILP ensures that, out of a set of domains *C* that all conflict with each other, only one is chosen 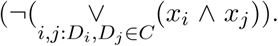 In practice, the cliques *C* above are found with a greedy clique-finding heuristic, returning *C* that are not nested but may overlap, and cover all conflicting domain pairs.

**Figure 1:**
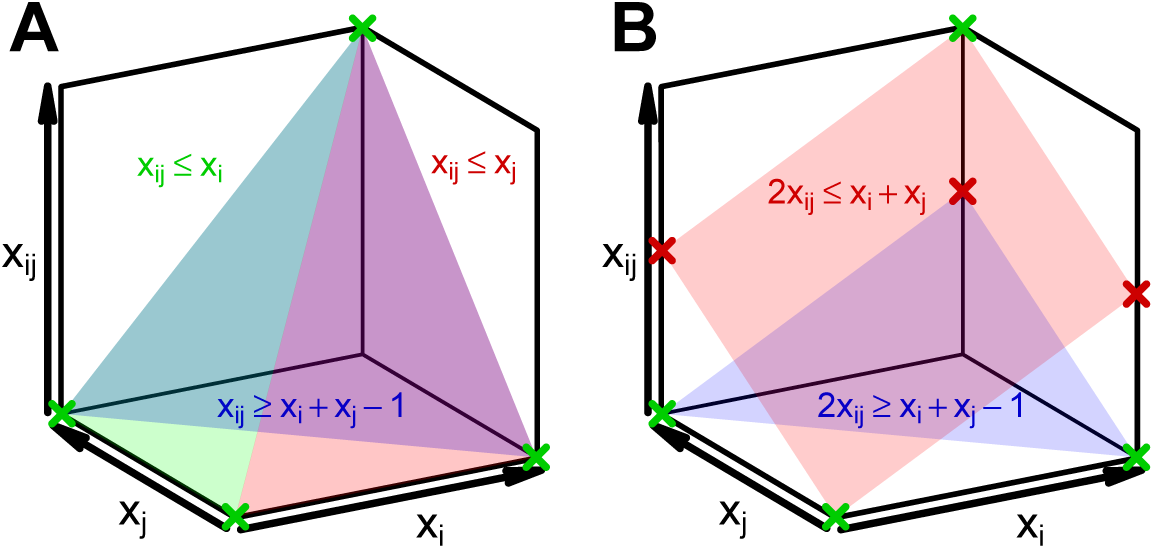
**A.** Geometric view of the relaxed LP constraints of Eq.(3)-(Eq.(5)). All *x_i_*, *x_j_* and *x_i_,_j_* are constrained by Eq.(2) to the unit cube shown. Equation colors match their translucent surfaces, and the blue surface is behind the others. The vertices of the polytope resulting from the intersection of these three constraints and the cube arise at integral values of *x_i_*, *x_j_* and *x_i_,_j_* (green crosses). **B.** Previously, we described another ILP [6] that consisted of two constraints relating *x_i_*, *x_j_*, and *x_i_,_j_*; that formulation is weaker than our current one as its polytope (the space between the two planes and inside the unit cube) is larger and has undesirable fractional vertices (red crosses) along with integral vertices (green crosses).

#### 3.3 Evaluation and comparison to other methods

##### 3.3.1 Pfam database

All tests use Pfam 25 (12,273 HMMs). Pfam provides curated thresholds, clan definitions (which group related families), and a list of nesting families that may overlap without conflicting. Family pair counts *c_i_,_j_* are observed on UniProt 2010_05 (11,384,036 proteins), as provided in Pfam-A.full. For dPUC2, Pfam sequence and family thresholds are used, with the exception that for simplicity, if 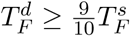(67 families), we set 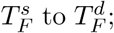 we note that this leaves only 163 families where sequence and domain thresholds differ.

### 3.3.2 Hmmscan parameters

Domains are predicted with `hmmscan` from HMMER 3.0. We used: “`--F1 1e-1--F2 1e-1 --F3 1e-2`” to set “stage 1/2/3” p-value thresholds and “`-Z 1 --domZ 1`” to return *p*-values instead of *E*-values. For testing our approach against others and computing false discovery rates (FDRs) across a range of noise levels, we used “`-E 1e-2 --domE 1e-2`” to set a permissive *p*-value threshold. For measurements of runtime, we used “`--cpu 0` ” to use a single thread (no parallel worker threads), and “`-E 1e-4 --domE 1e-4`” to set final *p*-value thresholds, as this is how our tool dPUC2 would be used in practice.

### 3.3.3 FDR Tests

We previously introduced five domain prediction FDR tests based on different definitions of true and false positives (TP, FP) [11]. Briefly, each FDR test labels predictions as TPs or FPs (as described below). Let TP*_i_* and FP*_i_* be the number of true and false positive labels predicted for protein *i*, respectively. The estimated FDR across *m* proteins with domain predictions is

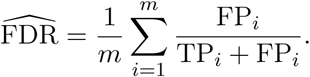

RevSeq (Reverse Sequence) pools domain predictions from real and reversed UniRef50 sequences (version 2011_04, 3,865,311 proteins). Different methods select domains from this set, treating all domains as belonging to one sequence (Fig. 2A). Selected domains on real sequences are labeled as TPs, and those on reversed sequences are labeled as FPs. To yield sensible FDRs, reverse sequence domains that overlap real sequence domains of the same family are removed (as some domain families have palindromic signatures), and the 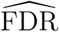 is doubled [11].

**Figure 2:**
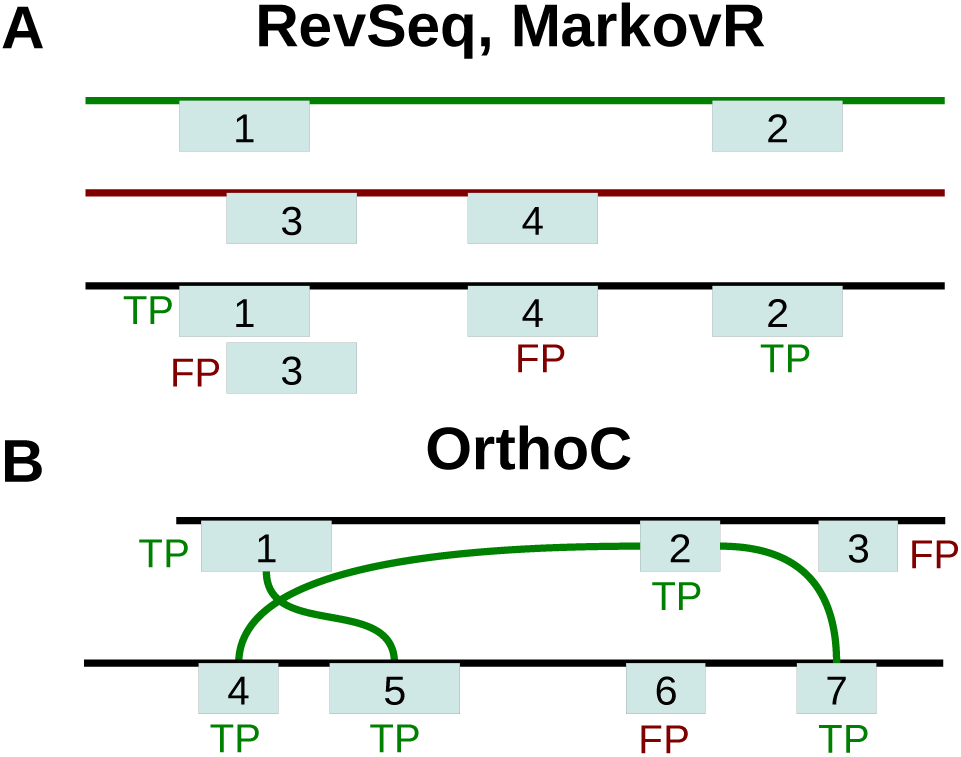
Illustration of FDR tests. **A.** RevSeq and MarkovR have a real (green line) and random sequence of the same length (reversed or Markov sequence; red line). Methods select domains pooled from both sequences (black line), and real sequence domains (boxes 1, 2) are TPs, while random sequence domains (boxes 3, 4) are FPs. **B.** OrthoC labels domains as TPs if there are domains of the same clan with *p* <1e-4 in orthologs (connected by green edges: boxes 1, 5; 2, 4, 7), FPs otherwise (the clans of 3, 6 are not in orthologs).

MarkovR (Markov Random) generates random sequences from a 2nd-order Markov model of UniRef50 [11], implemented by RandProt (http://github.com/alexviiia/RandProt). Real and random sequence domains are merged, and then MarkovR FDRs are computed similarly to the RevSeq FDR test (Fig. 2A).

Lastly, OrthoC (Ortholog Set Coherence) labels domains in a sequence as TP or FP if domains of the same clan are present in its orthologs or not, respectively (Fig. 2B), as described earlier [11]. Orthologs from OrthoMCL5 [14] are processed as before [11], with extra filters to eliminate proteins that do not share the same domain architecture as other proteins in their ortholog group. In particular, ortholog groups are now filtered so that sequences are iteratively removed if (1) they exceed the median length of their ortholog group by 50 amino acids (as they may contain additional domains), and (2) if they contain domains with HMMER *p*-value <1e-10 that are not found at the clan level in any of its orthologs with *p*-value < 1e-4 (as these predictions are high confidence and should not be considered FPs even if other sequences in the same ortholog group do not contain instances). Further, each ortholog group is pruned so that no two sequences share >90% identity. This results in a total of 79,386 ortholog families that consist of 525,258 sequences. We note that the two other previously introduced tests [11], ContextC and ClanOv, cannot be used to evaluate dPUC2, since they use context and require predicted domains with conflicts, respectively.

### 3.3.4 Comparisons to other approaches

We first compare how well our approach identifies domains as compared to Pfam when using several different approaches for setting thresholds. The “Standard Pfam” predicts domains using the Pfam domain and sequence thresholds, but removing domain instances of the same clan that overlap strictly (one or more amino acids), keeping only the most significant prediction per conflict [12]. To get a range of FDRs using Pfam, we “extend” the Standard Pfam thresholds by shifting domain and sequence thresholds by constant bit amounts [6]. As an alternative approach, we also consider a version of Pfam where predictions are considered as *E*-value thresholds are varied. Both Extended Standard Pfam and *E*-values remove conflicting domains (defined as permissive overlaps, see Models) ranking by *p*-value.

We also compare the performance of our method to three previous methods that use domain context. First, DAMA [5] was downloaded from http://www.lcqb.upmc.fr/DAMA and used with default Pfam 25 inputs. DAMA combines *E*-values with discrete rewards for previously-observed domains architectures and other criteria [5]. Next, we reimplemented the rule-based system CODD [4, 7], as a version using Pfam 25 and UniProt is not publicly available. CODD uses a list of “certified domain pairs”, a filtered binary context network, to predict low-confidence domains that are supported by context with high-confidence domains [4]. For CODD, certified domain pairs are defined [4] as Pfam 25 domain pairs occurring in UniProt that significantly co-occur (*p* < 0.01 using the Hypergeometric distribution). Lastly, we also test our previous dPUC1 approach, which does not take into account directional context. In this case, due to runtime limitations, it is not feasible to run dPUC1 directly in large-scale testing; instead, the dPUC1 objective is solved using the dPUC2 ILP outlined here. We did not test the method of [3], as it is not publicly available.

To compare the context methods, we first use a permissive *p*-value threshold of *p* < 1e-2 to get a set of candidate domains for each protein sequence. Next, we vary the *p*-value threshold between 1e-9 and 1e-3; in total, 17 thresholds are sampled (1e-9, 2e-9, 4e-9, 6e-9, 1e-8, 2e-8, …, 6e-4, 1e-3). We run DAMA with these domain predictions and each *p*-value threshold. For dPUC2, dPUC1 and CODD, the input is the set domains that are predicted by HMMER at each threshold, along with all Standard Pfam domain predictions. Finally, we calculate FDRs for each method using each of the three tests, and consider as a function of FDR, the change in number of domains as compared to the Standard Pfam.

## 4 Results

### 4.1 DPUC2 outperforms competitors in FDR tests

We compare the performance of dPUC2, our tool that implements the probabilistic directional domain context approach described in this paper, to five other approaches for domain identification. We begin by observing that across all three tests, context methods improve substantially over the Standard Pfam (Fig. 3). For example, dPUC2 predicts roughly 15% more domains than the Standard Pfam at the same empirical FDR. We note that all context methods improve upon the uncurated version of Pfam, where *E*-value thresholds are simply varied, across the entire range of estimated FDRs. In practice, however, Pfam uses curated domain and sequence thresholds; if these values are varied instead, the context method DAMA is outperformed by Pfam at low FDRs in two of our three tests.

**Figure 3:**
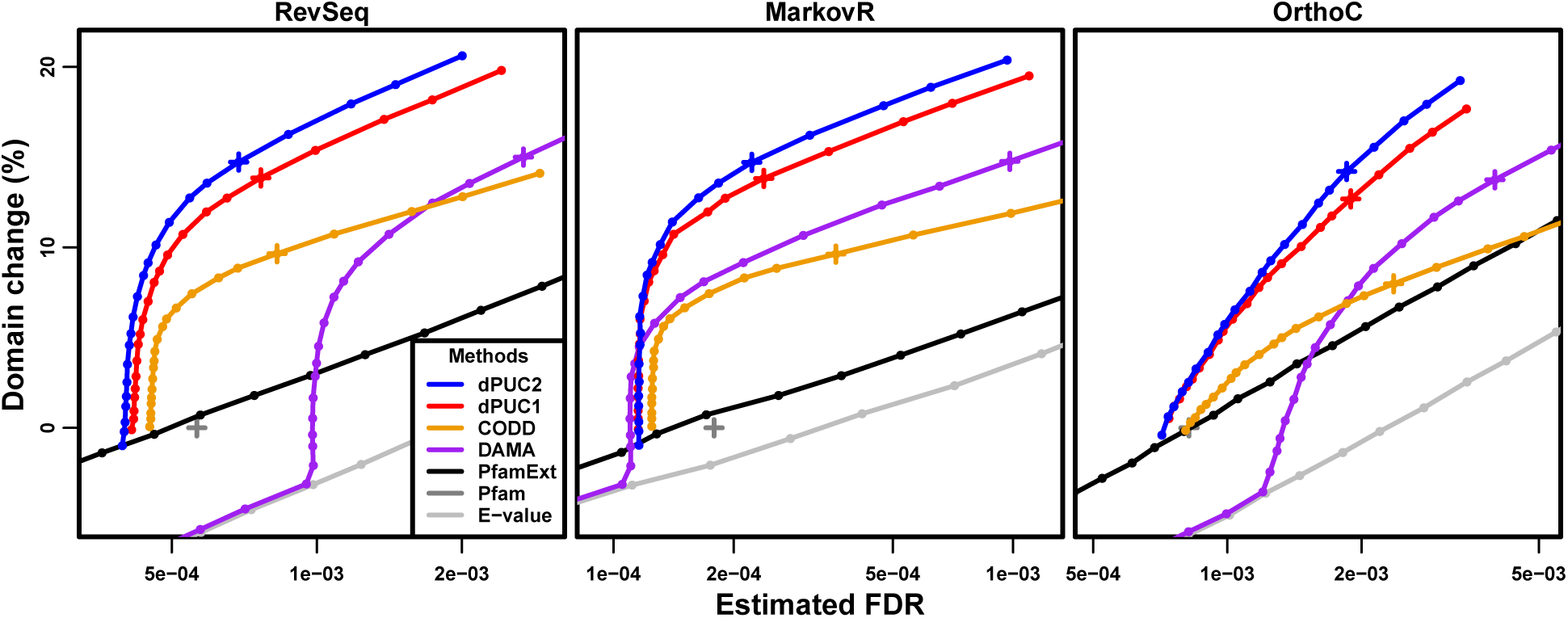
Our new approach, dPUC2, predicts more domains than its competitors across a wide range of FDRs, as estimated by the RevSeq (left), MarkovR (middle), and OrthoC (right) tests. The dark gray cross gives the FDR of the Standard Pfam, and the changes in the number of domain predictions for all methods at different FDRs are given with respect to the number of Standard Pfam predictions. For reference, we highlight with crosses the performances of dPUC2, dPUC1, CODD and DAMA when run on candidate domains identified with HMMER at *p* < 1e-4.

We find that dPUC2 outperforms the previous context-based methods in each of the three tests (Fig. 3). Notably, in all three tests, dPUC2 predicts more domains than CODD across the entire range of estimated FDRs. CODD does not penalize domain combinations that have not been observed before (i.e., it has no negative context) and this contributes to its higher FDRs. CODD also predicts many fewer domains, in part because no domains can be predicted by CODD in sequences that lack Standard Pfam predictions [4]. Our approach dPUC2 also consistently predicts more domains across the range of FDRs than the context method DAMA in the RevSeq and OrthoC tests. In the MarkovR test, when very low *p*-value thresholds are used to identify candidate domains (*p*-value < 6e-7), dPUC2 identifies more domains than DAMA but also results in higher estimated FDRs; this is due in part to dPUC2 keeping all Standard Pfam predictions in its input (see below). Thus, while overall dPUC2 performs better than DAMA, when using highly confident domain predictions, dPUC2 and DAMA trade off coverage and noise in the MarkovR test. Finally, in every FDR test, dPUC2 also improves upon dPUC1, illustrating the benefit of incorporating directional context. Directional context reduces the FDR of predictions produced for the same inputs; for example, when using candidate domains with *p* < 1e-4, dPUC2 results in a lower FDR than dPUC1 (blue and red crosses in Fig. 3).

We note that the dPUC2, dPUC1, and CODD methods always keep the Standard Pfam predictions in the input, so the candidate *p*-value threshold is applied to the non-Standard Pfam predictions only. For this reason, as more stringent thresholds are used to identify candidate domains, these methods perform similarly to the Standard Pfam predictions (dark gray cross in Fig. 3), differing only by how they remove predictions using negative context. On the other hand, for stringent *p*-value thresholds, DAMA removes those Standard Pfam predictions that are not predicted at that level of confidence. Therefore, DAMA performs similarly to the *E*-value curve when the *p*-value threshold is stringent, and differences between DAMA and the other context methods at low *p*-value thresholds are affected by the differences in the way the approaches handle Standard Pfam predictions. However, DAMA also incurs high FDRs at larger *p*-value thresholds which would include Standard Pfam predictions; for example, when using *p*< 1e-4 to identify candidate domains, DAMA’s FDR is higher than those for the other context methods in all three tests (purple cross in Fig. 3). Thus, the superior overall performance of dPUC2 as compared to DAMA is due to a variety of methodological factors.

### 4.2 Accelerated performance

DPUC2 is optimized to be much faster than dPUC1. We compare the dPUC1 and dPUC2 runtime using candidate Pfam domain predictions with *p* < 1e-4 on the *Homo sapiens* proteome (UniProt UP000005640, dated 2015-06-07, 69,693 proteins). Using a 3.2 GHz processor with 16 GB RAM, we find that the HMMER3 step that predicts candidate domains averages 0.455 second per protein (s/protein) in wallclock runtime when run in a single thread; this is comparable to a previous reported average of ~1 s/protein [2]. The dPUC2 step (excluding the HMMER3 runtime) averages 5.5e-3 s/protein, while dPUC1 averages 0.291 s/protein (Fig. 4). Therefore, dPUC2 is 53 times faster than dPUC1 on average, and amounts to 1.21% of HMMER3 runtime. Since the runtime distributions are highly skewed, we also compare median runtimes: the dPUC2 median is 1.11e-4 s/protein, while the dPUC1 median is 0.0282 s/protein; that is, the dPUC2 median run is 255 faster than dPUC1’s. Additionally, dPUC2 is faster than dPUC1 in 99.2% of individual proteins. Hence, dPUC2 is orders of magnitude faster than dPUC1 and its runtime is negligible compared to that of initially identifying candidate domains via HMMER3.

**Figure 4:**
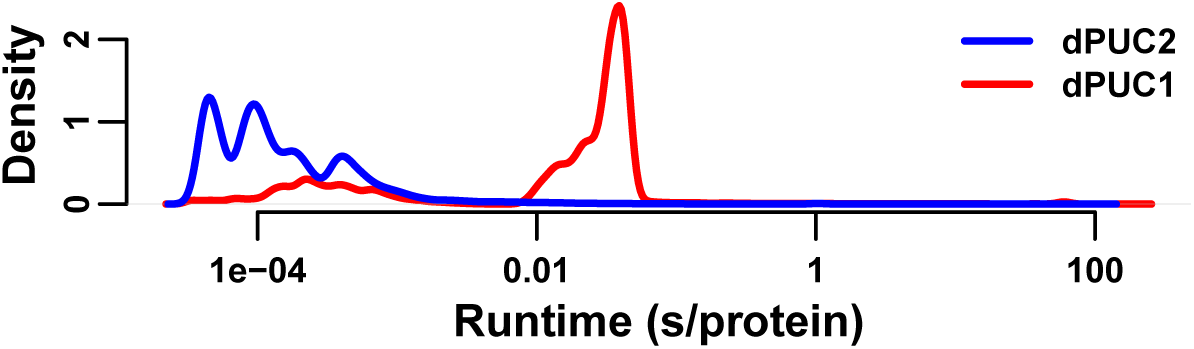
Distribution of wallclock runtimes on human protein sequences using dPUC1 or dPUC2 on a 3.2 GHz processor. A candidate domain threshold of *p* < 1e-4 was used in both cases. Density is for the log of the runtime.

## 5 Discussion

In this work, we introduce dPUC2, the first domain context prediction method that models directional domain context between all domain pairs, and perform the largest-scale evaluation to date of domain context methods. Using large and unbiased protein sets and several different criteria to estimate the FDR, we demonstrate that dPUC2 substantially outperforms previous methods that do not model context directionally or probabilistically (Fig3. 3). We note that previous evaluations of context-based domain identification approaches focused on the *Plasmodium falciparum* proteome [4–7]; however, our FDR tests—based upon the diverse UniRef50 database—are a better reflection of the true performance of context-based methods on proteins across all organisms.

Several features of dPUC2 are likely to contribute to its performance benefits over other currently available methods. One major advantage of dPUC2 over the previous context-based methods CODD and DAMA is that dPUC2 models context using domain pair-specific probabilities. In contrast, CODD rewards all “conditionally dependent pairs” equally, regardless of their frequency. Similarly, DAMA assigns the same rewards to all previously-observed domain architectures, ignoring their frequency. Additionally, neither CODD nor DAMA penalize domain combinations that have not been observed before whereas dPUC2 identifies domains by balancing scores arising from HMM matches with those arising from domain pairings. As compared to dPUC1, dPUC2 considers the order in which domains occur in protein sequences and additionally is more than 50 times as fast; the superior performance of dPUC2 illustrates the importance of modeling directional context. Finally, our newly introduced optimization techniques have resulted in runtimes for dPUC2 that are negligible compared to those of running HMMER3; thus, it is feasible to routinely consider context when making domain predictions.

In the future, it is likely that context-based methods could be further improved by using *q*-values [15] instead of *p*-values or *E*-values for candidate domain identification; though HMMER3 does not yet use *q*-value thresholds for domain identification, recent work has demonstrated that they lead to superior performance [11]. Another important line of future work for context-based methods is that of assigning a confidence to each individual domain prediction after considering context; one possibility would be to update the HMMER3 domain *p*-values given context or otherwise estimating posterior error probabilities. More generally, while we and others have shown the power of considering domain co-occurrence for domain identification, a deeper statistical understanding of domain prediction using context is likely to lead to still better methodologies.

## Acknowledgments

We thank Manuel Llinás for helpful discussions about this work.

## Funding

This work has been supported by the National Science Foundation [Graduate Research Fellowship DGE 0646086 to AO, ABI 1062371 to MS]; and the National Institutes of Health [R21-AI085415 and R01-GM076275 to MS].

## References

[1] S. R. Eddy. “A Probabilistic Model of Local Sequence Alignment That Simplifies Statistical Significance Estimation”. PLoS Comput Biol 4(5) (2008), e1000069.

[2] R. D. Finn et al. “The Pfam protein families database”. Nucl Acids Res 38(suppl_1) (2010), pp. D211–222.

[3] L. Coin, A. Bateman, and R. Durbin. “Enhanced protein domain discovery by using language modeling techniques from speech recognition”. P Natl Acad Sci U S A 100(8) (2003). PMC404693, pp. 4516–4520.

[4] N. Terrapon et al. “Detection of new protein domains using co-occurrence: application to Plasmodium falciparum”. Bioinformatics 25(23) (2009), pp. 3077–3083.

[5] J. S. Bernardes et al. “A multi-objective optimisation approach accurately resolves protein domain architectures”. Bioinformatics (2015), btv582.

[6] A. Ochoa, M. Llinás, and M. Singh. “Using context to improve protein domain identification”. BMC Bioinformatics 12(1) (2011), p. 90.

[7] A. Ghouila et al. “Identification of Divergent Protein Domains by Combining HMM-HMM Comparisons and Co-Occurrence Detection”. PLOS ONE 9(6) (2014), e95275.

[8] G. Apic, J. Gough, and S. A. Teichmann. “Domain combinations in archaeal, eubacterial and eukaryotic proteomes”. J Mol Biol 310(2) (2001), pp. 311–325.

[9] T. U. Consortium. “Reorganizing the protein space at the Universal Protein Resource (UniProt)”. Nucleic Acids Research (2011).

[10] C. Yeats, O. C. Redfern, and C. Orengo. “A fast and automated solution for accurately resolving protein domain architectures”. Bioinformatics 26(6) (2010), pp. 745–751.

[11] A. Ochoa et al. “Beyond the E-Value: Stratified Statistics for Protein Domain Prediction”. PLoS Comput Biol 11(11) (2015), e1004509.

[12] M. Punta et al. “The Pfam protein families database”. Nucleic Acids Research 40(D1) (2011), pp. D290–D301.

[13] M. Berkelaar, K. Eikland, and P. Notebaert. lp_solve: Open source (Mixed-Integer) Linear Programming system. 2004.

[14] F. Chen et al. “OrthoMCL-DB: querying a comprehensive multi-species collection of ortholog groups”. Nucleic Acids Res. 34(Database issue) (2006). PMC1347485, pp. D363–D368.

[15] J. D. Storey. “The positive false discovery rate: a Bayesian interpretation and the q-value”. Ann. Statist. 31(6) (2003). Mathematical Reviews number (MathSciNet): MR2036398; Zentralblatt MATH identifier: 02067675, pp. 2013–2035.

